# Nipoppy: A framework for standardizing neuroimaging studies to facilitate international derived-data sharing

**DOI:** 10.64898/2026.05.18.723593

**Authors:** Nikhil Bhagwat, Michelle Wang, Mathieu Dugré, Julia-Katharina Pfarr, Alyssa Dai, Sebastian Urchs, Brent McPherson, Rémi Gau, Eva M. van Heese, Emile d’Angremont, Max A. Laansma, Shweta Prasad, Jacob Sanz-Robinson, Mohammad Torabi, Arman Jahanpour, Matthew Danyluik, Alice Joubert, Austin Macdonald, Lea Waller, Ashley Stewart, Matthieu Joulot, Erin Dickie, Gabriel A. Devenyi, Sylvain Bouix, Steffen Bollmann, Neda Jahanshad, Paul M. Thompson, Ninon Burgos, M. Mallar Chakravarty, Yaroslav O. Halchenko, Ysbrand D. van der Werf, Jean-Baptiste Poline

**Affiliations:** McConnell Brain Imaging Centre, The Neuro (Montreal Neurological Institute-Hospital), McGill University, Montreal, Canada; Department of Computer-Science and Software Engineering, Gina Cody School of Engineering and Computer Science, Concordia University, Montreal, Canada; MIND team at Inria Saclay - Île-de-France; Department of Anatomy & Neurosciences, Amsterdam UMC, Vrije Universiteit Amsterdam, Amsterdam, the Netherlands; Amsterdam Neuroscience, Neurodegeneration, Amsterdam, the Netherlands; Department of Neurology, National Institute of Mental Health and Neurosciences (NIMHANS), Bengaluru, India; Cerebral Imaging Center, Douglas Research Center, McGill University, Verdun, QC, Canada; Department of Psychiatry, McGill University, Montreal, Canada; Sorbonne Université, Institut du Cerveau - Paris Brain Institute - ICM, CNRS, Inria, Inserm, AP-HP, Hôpital de la Pitié Salpêtrière, F-75013, Paris, France; Center for Open Neuroscience, Department of Psychological and Brain Sciences, Dartmouth College, Hanover, NH, USA; Department of Psychiatry and Neurosciences CCM, Charité Universitätsmedizin Berlin, corporate member of Freie Universität Berlin and Humboldt-Universität zu Berlin, Berlin, Germany; School of Electrical Imaging and Computer Science, University of Queensland, Australia; Centre for Addiction and Mental Health, University of Toronto, ON, Canada; Imaging Genetics Center, Stevens Neuroimaging and Informatics Institute, Keck School of Medicine, University of Southern California, Los Angeles, CA, USA; Département de génie logiciel et TI, École de technologie supérieure, Montréal, Canada; Department of Biomedical Engineering, McGill University, Montreal, Canada; German Center for Mental Health (DZPG), partner site Berlin-Potsdam, Berlin, Germany

## Abstract

Neuroimaging data management and processing are tedious and error-prone, prompting reproducibility concerns. Globally, studies with heterogeneous infrastructure and governance policies lead to eclectic data processing and sharing, necessitating standardization of data workflows to ensure reusability and comparability of multi-centric datasets. The Nipoppy neuroinformatics framework facilitates such standardization by combining specification, protocol, and software to manage study-level data workflows. With its adoption, researchers can share standardized, derived datasets enabling efficient, reproducible, and inclusive research.

## Main text

The history of neuroscience is inundated with both impactful discoveries and replication shortfalls that make a persuasive argument for scientific data-sharing at a global scale ^1–5^. Shared access to well-curated large, globally inclusive datasets is critical for the development, validation, and translation of novel biomarkers, prognostic tools in the clinic ^6^. At scale evaluations ascertaining reliability and generalizability of scientific outcomes are especially pertinent in the ongoing shift towards the collaborative human-AI research paradigm. Nonetheless, geographical separation and data governance policies often pose technical and privacy issues that complicate and constrain *en masse* integration and analysis of global datasets ^5^. In a typical neuroimaging study lifecycle, collected data objects undergo several sequential transformations before they are statistically analyzed and reported in the form of experimental findings (Fig 1A). Notably, after a sequence of these transforms, the risks of identifying personal information from the pseudonymized “derived data” diminish, opening up data sharing mechanisms ^7^. Unfortunately, these transforms concomitantly induce methodological variance that severely hinders interoperability and reuse of these derived data across studies ^8^.

**Figure 1.**
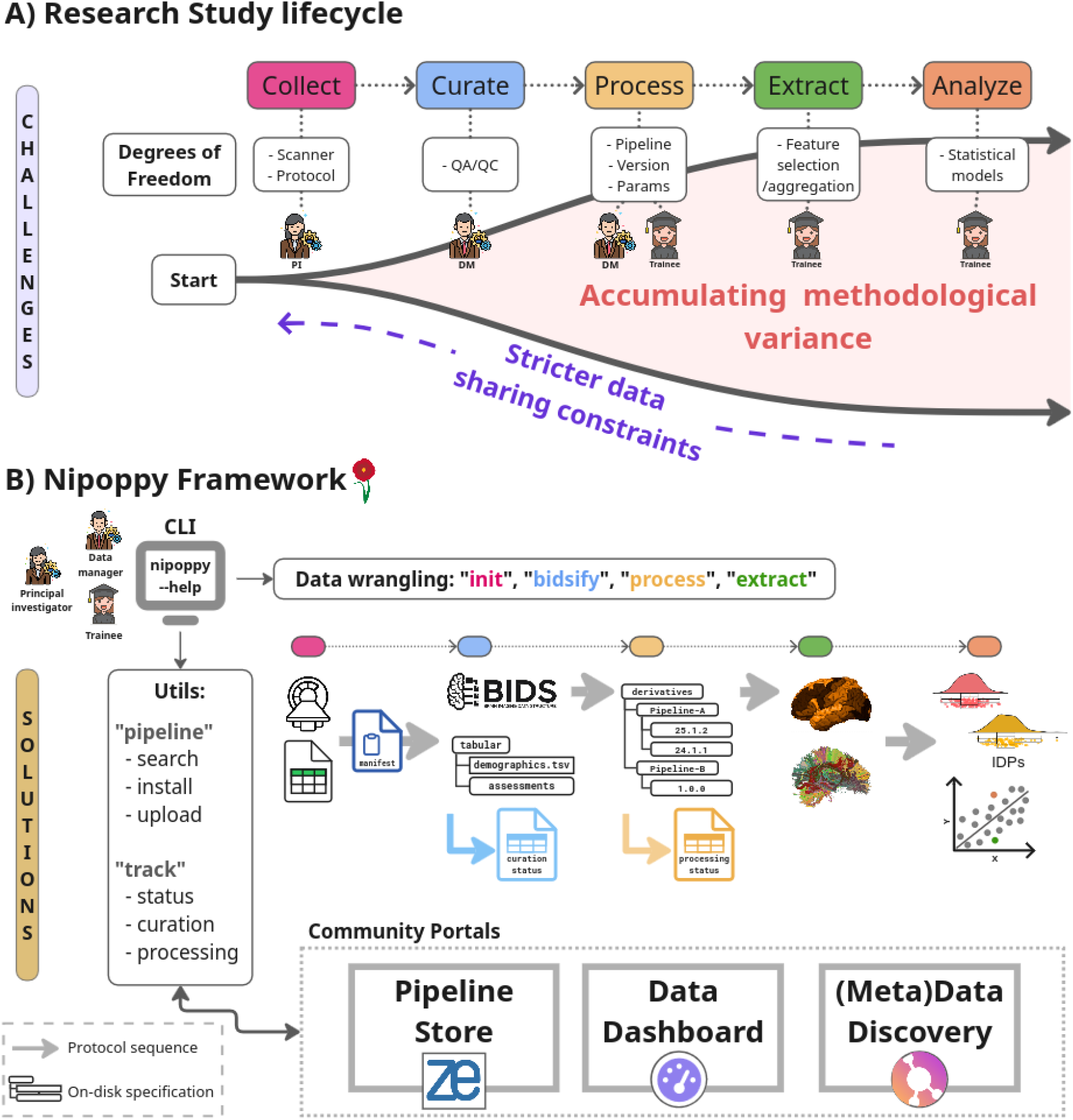
A) The conceptualized neuroimaging study cycle and associated challenges complicating data sharing and reuse. B) The Nipoppy solution to standardize data workflows at various study stages with adoption of a protocol, a specification, and a Python package to facilitate consistent data processing, discovery, and sharing.

Standardization of data workflows in a study lifecycle is an effective strategy for sharing consistent derived data in the community. The Nipoppy framework, based on the Findable, Accessible, Interoperable, and Reusable (FAIR) ^9^ principles and community standards, offers such a solution. It is unique in providing a triad of components: 1) a standard protocol for data workflows, 2) a specification for on-disk study-level data management, and 3) a Python package enabling generation of reproducible, analysis-ready derived datasets. In contrast with other solutions, Nipoppy’s functional, non-monolithic approach is a deliberate attempt to train and guide researchers through data wrangling tasks. Nipoppy does not interfere with existing data collection and ingests source data “as-is”. It supports neuroimaging and phenotype datatypes and is agnostic to scanner protocols and imaging pipelines. Nipoppy conceptualizes the data flow in a study into five sequential stages (Fig. 1): 1) Capture, 2) Curation, 3) Processing, 4) Extraction, 5) Analysis, providing tools and guidelines for each. This modular conceptualization can support a wide range of use cases while standardizing data organization and processing tasks, which has proved particularly effective in global consortia studies where consistent image processing across sites is crucial for mega and meta analyses ^10^.

Nipoppy is intended for human-in-the-loop data-wrangling scenarios and is guided by user-centric design principles. To promote and simplify adoption across a large spectrum of technical expertise, Nipoppy is developed to be *lightweight* for installation and usage while still allowing versatile use cases. The Python package comes with minimal dependencies and provides utilities (Fig 1B) to handle tedious technical tasks including pipeline installation and study status tracking. It helps users and study coordinators build a “dry lab protocol” (akin to wet lab protocols) specifying the *ordering of tasks* to standardize data workflows across multiple studies and sites. Nipoppy expects wrangling tasks to be *iterative*, and hence is built to simplify reproducibility, reuse, and replication of data workflows.

### A protocol

Nipoppy recommends an ordered sequence of steps, mapped onto the conceptualized study stages, with verifiable start and end points. This is particularly useful in neuroimaging studies where different tasks (e.g. imaging vs clinical data acquisition) and study stages (e.g. data curation vs processing) are handled by different people. The ordered sequence of steps operates on top of a specification that helps formalize and track expected outcomes in the study at various stages (see Methods). The protocol minimizes idiosyncratic deviations in decentralized setups and helps catch errors earlier in the study cycle.

### A specification

Nipoppy directory layout and file schema extend the Brain Imaging Data Structure (BIDS) standard ^11^ to support data management of source (i.e. collected), raw (i.e. standardized), and derived (i.e. processed) data objects generated during a study (Fig S1). Leveraging Boutiques^12^, the specification allows users to standardize curation, processing and tracking of on-disk datasets during the entire lifecycle of a study comprising both imaging and phenotypic modalities. The specification is agnostic to imaging modalities and processing pipelines which can be rapidly shared and deployed using the built-in Zenodo interface.

### The Python package

A command-line interface (CLI) operationalizes the Nipoppy protocol and specification. The CLI provides generalized commands to run virtually any pipeline for data curation, processing and extraction tasks. The helper utility commands allow users to track pipeline outputs and share pipeline-bundles across studies and sites. The `nipoppy` package drastically simplifies iterations of relaunching pipelines on new or failed participants. The package supports high-performance computing job schedulers (Slurm, SGE, etc.) further simplifying launching pipelines at scale. The extensive documentation with guides, video tutorials and a Discord user forum helps lower the barrier for adoption and build a user and developer community invested in FAIR practices and data sharing.

The ease-of-use and immediate practical benefits have driven Nipoppy’s adoption. The current active users include over 50 international sites (Fig. 2A) with datasets comprising 10K+ participants from healthy and disease cohorts. The majority of sites come from consortia studies which often struggle with coordination of data processing and periodic pipeline updates resulting in inconsistent derived data (Fig. 2B). The ENIGMA Parkinson’s Disease working group (PD-WG) is the largest international neuroimaging effort in PD focusing on large-scale mega-analysis on multisite derived data. In late 2024, the PD-WG adopted the framework to standardize data management and processing across their sites, which rapidly improved data sharing efficiency and derived data consistency. By mid-April 2025, 15 sites had successfully processed their data from FreeSurfer 7, a task that took multiple years for the previous FreeSurfer version ^10^. Since then, PD-WG projects created standardized “pipeline-bundles” on Zenodo ^13^ (e.g. fMRIPrep ^14^, FreeSurfer_subseg ^15^, QSIPrep ^16^) which are seamlessly disseminated, run, and tracked at multiple sites (Fig 2C).

**Figure 2.**
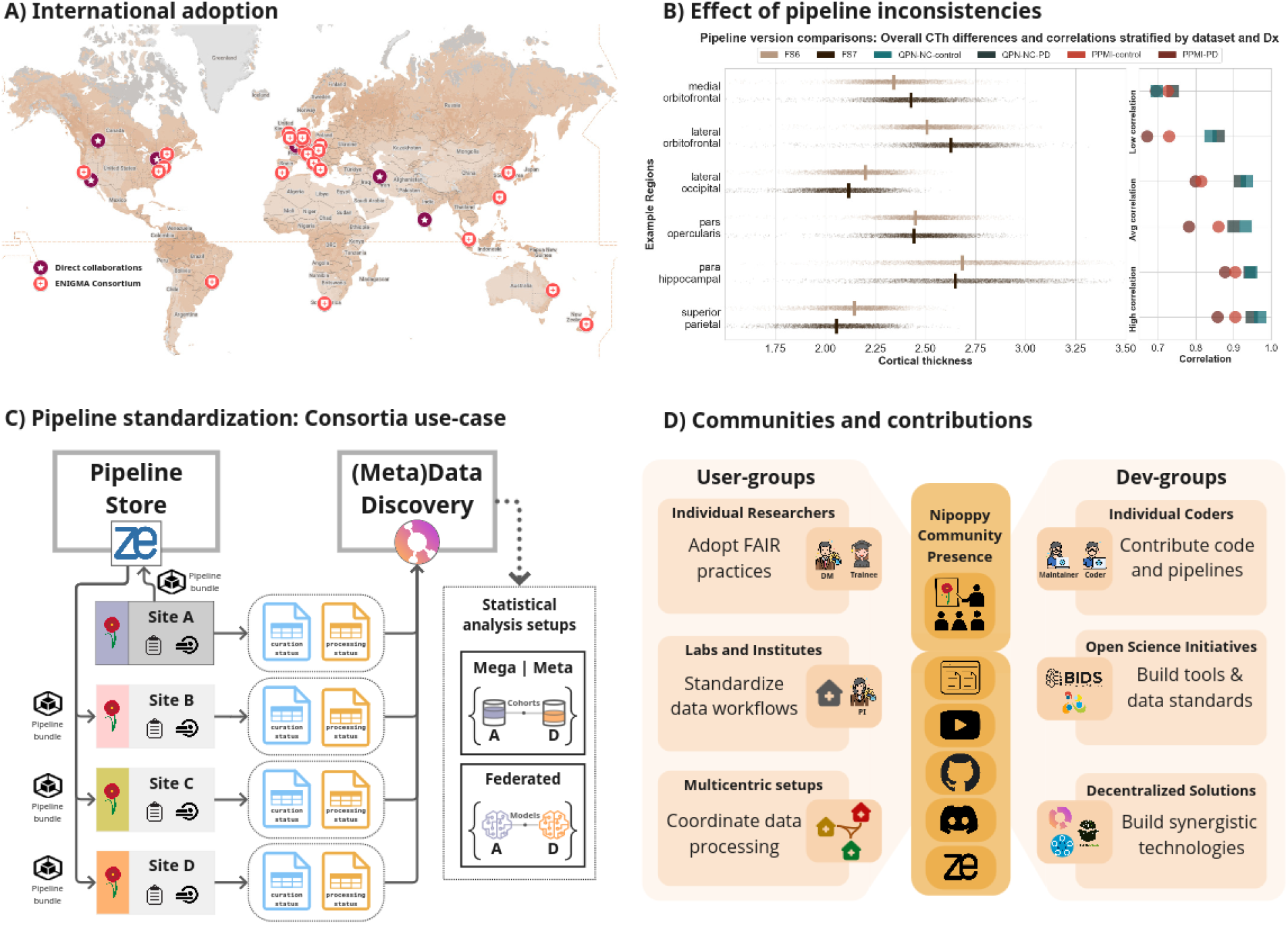
A) Nipoppy adoption by international research labs and consortia. B) Variability in cortical thickness (CTh) estimation between two FreeSurfer versions compared in two Parkinson’s disease datasets. See supplement for more details. C) Standardized deployment of custom pipelines followed by tracking, discovery, and sharing of derived data for analysis in a consortia setup. See “Use case” section in the Methods. D) Nipoppy community building efforts and open-science contributions.

Nipoppy contributes to existing community standards and supports integration with several related informatics initiatives (Fig 2D). Primarily, it builds on the BIDS-study specification, with added files and directives that facilitate the “protocol” aspect, while maintaining full compatibility. The Nipoppy developer team also maintains the Boutiques package used to launch pipelines and record their metadata and runtime parameters. Additionally, Nipoppy is expanding its support to a broader range of neuroimaging pipelines by integrating with the Neurodesk ^17^ platform, automating derived data discovery and sharing via Neurobagel ^18^, and serving analysis-ready derived dataset for federated learning with Fed-BioMed ^19^.

The focus and design choices of Nipoppy driven by its scope and user-base include certain functional tradeoffs. Nipoppy primarily supports neuroimaging datatypes, with minimal specification for the management of phenotypic (i.e. demographics, clinical-assessments) data. It is not a workflow engine and therefore cannot support workflows with complex pipeline dependencies and automated executions. While it tracks processing in terms of pipeline versions and run-time parameters, to remain lightweight, it does not capture file modifications. Lastly, management of Zenodo hosted pipeline-store incurs curation burden. Although many potential containerized pipeline hubs can be used with Nipoppy, only a modest number of vetted pipelines remain directly searchable in the Nipoppy community. Such scaling challenges are being addressed through a community governance model to ensure long-term sustainability of the project.

The multidimensional technical, governance, and social challenges associated with global data sharing requires a multifaceted solution. Nipoppy uniquely combines protocol, specification, software, and training efforts into a single, user-friendly framework founded on FAIR practices that helps researchers to conceptualize and navigate complex data-wrangling and re-use. Nipoppy is growing as a study level standard and an extended BIDS community initiative to lower the barriers for collaborations across global institutes, scale-up data sharing, and create a more inclusive data-future for neuroimaging research.

## Methods

### Nipoppy Terminology

Glossary of important terms used in the Nipoppy framework:

1. Study: A human-in-the-loop process involving multimodal measurements from a set of participants to be processed for scientific analysis.
2. Workflow: A sequential set of manual and automated tasks involving data curation, processing, extraction, and analysis operations.
3. Pipeline: A software package to transform data with the purpose of standardization, image processing, feature extraction, or statistical analysis.
4. Collection: A task of capturing data points from a participant prior to any transformations.
5. Curation: A task of transforming collected data into a community standard (e.g. BIDS) and / or user-friendly data format (NIfTI).
6. Processing: A task that takes the curated data as input and applies an image processing pipeline (e.g. image registration, segmentations) to generate a new set of images.
7. Extraction: A task that takes the processed data as input and applies feature extraction pipelines to derive a set of phenotypes (e.g. regional volumes, tract-specific diffusion measures, brain atlas associated networks) that can be directly used towards statistical analysis.
8. Tracking: A task that checks the presence of a set of files on the disk.

### How to Nipoppify your study: Protocol adoption in practice

The Nipoppy framework can be adopted by retrospective, on-going, or prospective studies and is designed to accommodate human-in-the-loop tasks. In practice, the user – typically a data manager or a researcher – utilizes the Python package in conjunction with manual tasks to follow the Nipoppy protocol. The protocol begins with initialization of an on-disk directory tree layout as per the specification. The various files and directories in this layout are populated as the user progresses through the study stages. Next, the user generates the `manifest.tsv` comprising a list of participants and their visits in the study. The manifest serves as the interface between the data capture and curate stages and contains *expected availability* information based on which downstream tasks are run and tracked. The entries in the manifest can be gathered from multiple data capture sources (e.g. DICOM server, clinical databases, electronic health records) automatically, but often require manual validation to confirm updated study enrollments. The modality agnostic manifest simplifies iterative execution and tracking of all data workflows prompted by cohort additions and pipeline upgrades.

Subsequently, based on a study design, the user installs a set of pipeline-bundles (containers and associated configuration files) for data curation, processing, and extraction workflows. Users can easily search and install commonly used neuroimaging pipelines from the public Zenodo repository maintained by the Nipoppy team or, alternatively, they can create and use custom pipelines locally. Pipelines can be installed and updated at any step of the protocol.

In the next step of the protocol – data curation – the user transforms the source data for the participants listed in the manifest into the BIDS standard. The Nipoppy CLI provides helper functions to reorganize source data to simplify the actual BIDSification process with heuristic components. The generalized `bidsify` command allows users to work with virtually any BIDSification pipeline of choice. The data curation workflow finishes with the user verifying successful on-disk presence of raw-bids dataset within the `rawbids` directory by running a curation tracker in the CLI to generate the `curation_status.tsv` file. For behavioural and clinical tabulated data, Nipoppy supports BIDS phenotype extension and provides additional recommendations for organizing demographic variables.

The next step – data processing – is typically the most resource intensive and difficult neuroimaging task to manage at scale. Nipoppy’s generalized CLI commands significantly simplify pipeline installation, execution, and tracking of the multiplicative multiverse of participants x visits x pipelines x versions. Nipoppy’s on-disk layout and extended Boutiques specification helps users automate several iterative tasks, including re-processing a subset of participants and visits, checking processing statuses, and adding or upgrading pipelines as necessary. Users also benefit from the provenance of pipeline metadata and – importantly – runtime parameters, which ensures reproducibility locally and facilitates replication on other datasets and sites. The data processing workflow finishes with the user verifying successful on-disk presence of processed data within the `derivatives` directory by running a processing tracker in the CLI to generate the `processing_status.tsv` file. This file can be interactively visualized in a web dashboard locally.

The processed output from the pipeline often needs to be further transformed into analysis-ready imaging-derived-phenotypes (IDPs). This – feature extraction – step tends to be specific to research questions and can rely on a multitude of biological priors. In order to maintain reproducibility, users can build, run, and share phenotype extraction pipelines to generate analysis-ready dataframes used to produce their statistical findings. This becomes critical in several multisite setups involving mega-, meta-, and federated-analysis where consistency of the shared IDPs and details on their extraction method and applied priors (e.g. atlases) is essential for statistical modeling and comparisons. Similar to data *processing pipelines*, the Nipoppy specification greatly simplifies at-scale deployment of *extraction pipelines*, minimizes methodological variance across decentralized sites, and improves interoperability and reusability of the multisite data.

The modularity of the protocol provides flexibility to the users to adopt the Nipoppy framework at any stage of their study and simplify downstream steps in the protocol. The lightweight Python package lowers the barrier for community adoption, which in turn facilitates at-scale sharing of derived datasets.

### Use case: Nipoppy adoption in a multisite setup

In studies with decentralized data governance (e.g. consortia), Nipoppy offers a lightweight standardized approach to coordinate data management, processing, and extraction tasks. By leveraging the Zenodo API and the pipeline-store, participating sites can seamlessly share, update, and deploy their pipeline-bundles—consisting of specific versions and runtime parameters—across the collaborative network. The step-wise protocol together with generalized pipeline execution via the CLI simplifies decentralized coordination of installation and runtime parameterization for both novel or custom pipelines. This significantly accelerates the image processing and IDP extraction tasks while maintaining reproducibility by reducing the communication ambiguity and local idiosyncratic variations that often compromise multi-centric studies.

This concrete use case has been part of several multi-site studies under a consortium governance structure. Notably, the ENIGMA Parkinson’s Disease working group (PD-WG) comprising 40+ cohorts, ENIGMA-Tremor working group with 10+ cohorts, Schizophrenia Canadian Neuroimaging Database (SCanD) initiative with 4 cohorts have adopted Nipoppy for streamlining and coordinating their internal study-cycles. In these studies, the prototypical steps with the Nipoppy adoption are as follows (see Figure 2C):

1. All participating sites organize their data using the Nipoppy specification and install the Nipoppy python package.
2. A project-lead site proposes an analytic project. For data processing and extraction, specifies existing pipelines-bundles or builds and publishes the custom pipeline-bundles (containers + config + parameters), on Zenodo using Nipoppy CLI. See example of ENIGMA-PD pipeline bundle for FreeSurfer_subseg (doi:10.5281/zenodo.15877956).
3. The project sites pull the pipeline-bundles and run the data processing and extraction using Nipoppy CLI.
4. The consistent derived data are then shared for mega- and meta-analyses to the project-lead site.

This repeatable procedure becomes increasingly efficient at supporting new sites, new pipelines, and their version upgrades. Several commonly used pipelines are maintained by the Nipoppy team under Zenodo community (https://zenodo.org/communities/nipoppy/records), and users and consortia are able to create and maintain their own communities or pipeline hubs as well.

### Lowering the barrier for FAIR research

The impact of Nipoppy in the global neuroimaging community is multifaceted. Quantitatively, as of March 2026, Nipoppy has been adopted by 50+ international sites to manage over 70+ datasets, alongside 700+ downloads of the top 10 Nipoppy Zenodo pipelines. Beyond the numbers, Nipoppy has been instrumental in promoting FAIR data practices and the adoption of the community BIDS standard among trainees. The “dry lab protocol” helps operationalize the use of core neuroimaging tools and pipelines, simplifying the management of large datasets with multiverses of derivative outputs. Through comprehensive documentation, video tutorials, and community forums (see Manuscript Figure 2D), the framework lowers the barrier to entry for high-quality, reproducible neuroimaging research across a wide spectrum of technical expertise. Ultimately, this lightweight and scalable solution empowers individual laboratories and research institutes, furthering the collective effort toward open, inclusive, and replicable science.

### Community focused development and long-term sustainability

Nipoppy is meant to be developed *by* and *for* the neuroscience community, especially given the interdependent software and social aspects involved in such neuroinformatic initiative. It is currently supported by a community of contributors from over 10 laboratories. In bi-weekly open meetings, users and developers directly discuss different design perspectives, use-cases and improvement ideas. Nipoppy’s comprehensive documentation, (https://nipoppy.readthedocs.io/en/latest/) comprising how-to guides and tutorials, is actively kept up-to-date based on development progress and user feedback. All training content is openly available and used regularly in tutorials and workshops at multiple brainhacks and hackathon events by the core team and collaborators. There are active ongoing efforts to promote and increase compatibility with other tools in the neuroimaging software ecosystem. In particular, the processing status files produced by Nipoppy can be directly used by Neurobagel to discover the availability of pipeline outputs across studies. A Nipoppy how-to guide demonstrates DataLad coupling to achieve full data versioning and pipeline provenance for the studies. Additionally, increased support for various runtime setups enables seamless pipeline execution within a Neurodesk environment. All Nipoppy software and pipeline-bundles are built on existing well-supported community tools, and released open source with comprehensive documentation. The design principles minimize Nipoppy-specific dependencies for managing study lifecycle, promoting viable community usage without lock-in issues.

The Nipoppy layout is fully compatible with the BIDS-study specification, and the maintainer teams actively collaborate to streamline efforts towards community building and the long-term support of the standards. The dedicated funding for software development and maintenance is currently in the process of formalizing governance structure comprising multiple stakeholder labs. Over the past year, the close collaborations with the large international projects and consortia have prompted organic user-base growth. This increasing interest from both developer and user sides promises a sustainable community-driven future for Nipoppy.

## Acknowledgements

We would like to thank Qing (Vincent) Wang, Christopher J. Markiewicz, Tristan Glatard, Ines Gonzalez Pepe, and Ninad Aithal for their input on shaping and improving the Nipoppy framework and tools. We also note that the “persona” icons in the manuscript figures are designed using resources from Flaticon.com.

