## Supplement for "Nipoppy: A framework for standardizing neuroimaging studies to facilitate international derived-data sharing"

### Supplement: Example analyses to show importance of standardization of processing pipeline versions

We assessed the effect of pipeline versions - a common confound in the neuroimaging experiments - with a set of analyses. We used T1w magnetic resonance images from 1200 participants from two Parkinson's disease datasets: PPMI (n=1000) and QPN (n=200). The images were processed in Nipoppy using two FreeSurfer versions (6.0.1, 7.3.2) and the aggregated estimates of cortical thickness were calculated using the Desikan-Killiany (DK) atlas.

#### S1. Regional variation in cortical thickness estimation across pipeline versions

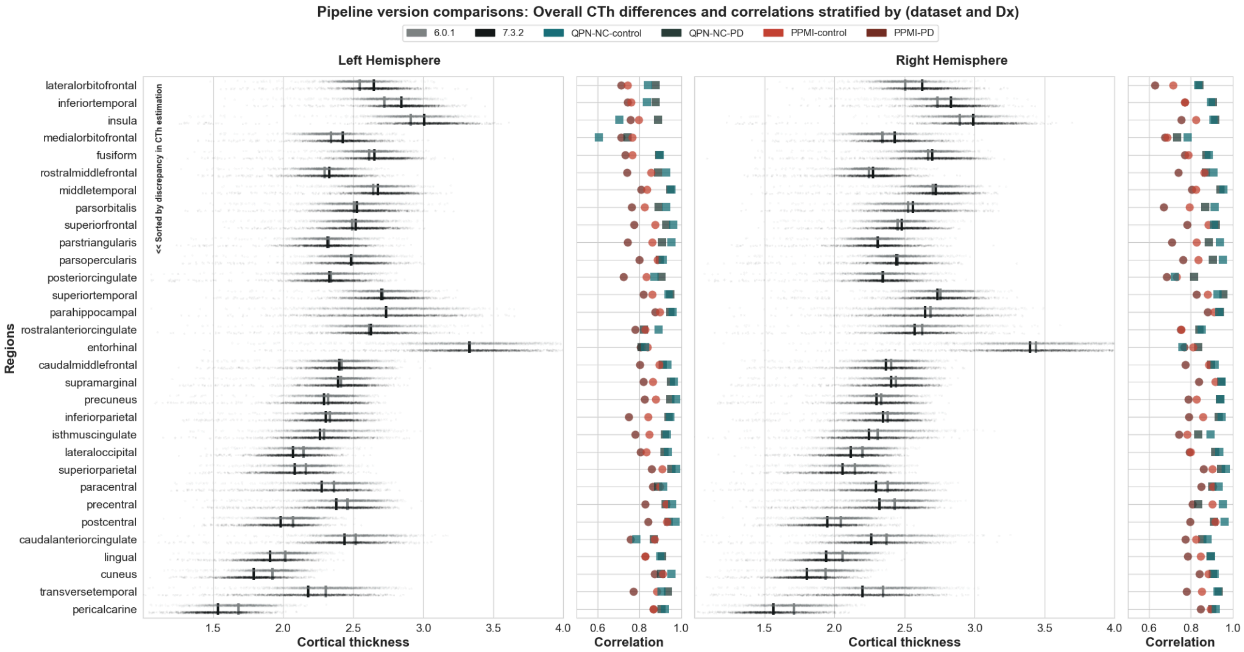


Figure S1. Variability in cortical thickness (CTh) estimation across the regions in the Desikan-Killiany (DK) atlas between two FreeSurfer versions. The scatterplot shows absolute CTh distribution differences between the two versions in an aggregate sample comprising two Parkinson's disease datasets. The pointplot shows correlation differences between versions stratified further by dataset and diagnosis. The differences illustrate the importance of standardizing pipeline versions.

### S2. Individual-level differences in cortical thickness estimation across pipeline versions

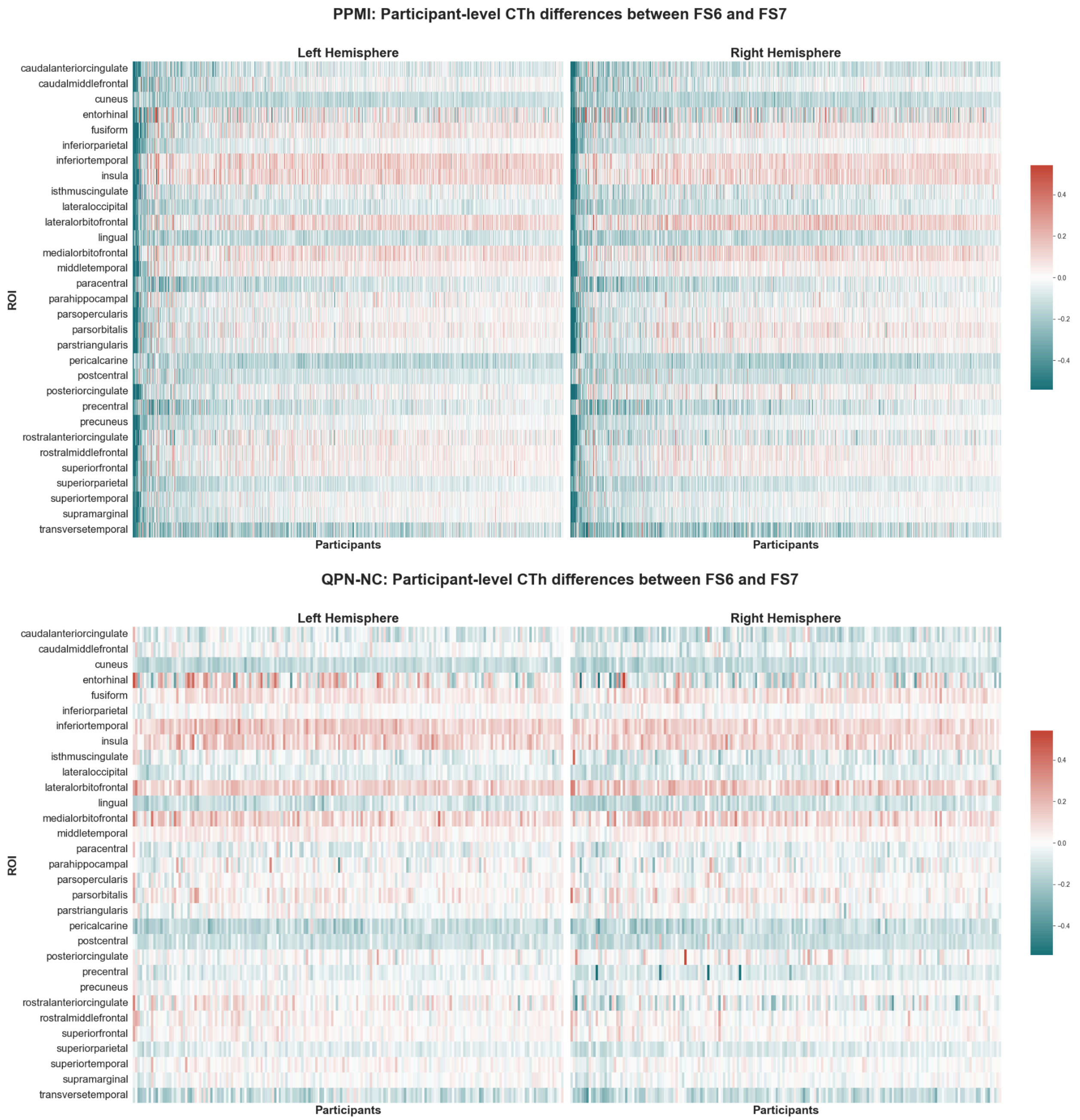


Figure S2. Heatmaps illustrating individual-level differences in CTh estimates between FreeSurfer versions across all Regions of Interest (ROIs). The results demonstrate high inter-participant variability with no apparent region-specific systematic shift, further complicating cross-study comparisons when pipeline versions are not identical.

**Differences in group-level differences (effect sizes) across pipeline versions**

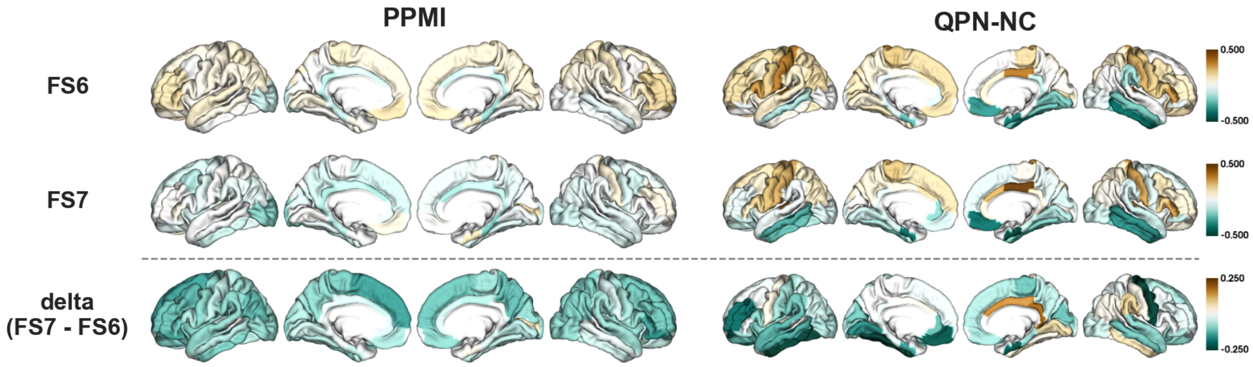


Figure S3. Spatial maps displaying Cohen’s d effect sizes for Parkinson’s disease (PD) vs. control group differences in CTh. The comparison across FreeSurfer versions and datasets shows that choice of pipeline version can significantly alter the spatial distribution and magnitude of reported effect sizes highlighting the importance of maintaining consistent processing pipelines.
